# A broad role for YBX1 in defining the small non-coding RNA composition of exosomes

**DOI:** 10.1101/160556

**Authors:** Matthew J. Shurtleff, Jun Yao, Yidan Qin, Ryan M. Nottingham, Morayma Temoche-Diaz, Randy Schekman, Alan M. Lambowitz

**Affiliations:** Department of Molecular and Cellular Biology, University of California, Berkeley 94720; Department of Plant and Microbial Biology, University of California, Berkeley 94720; Institute for Cellular and Molecular Biology and Department of Molecular Biosciences, University of Texas at Austin, Austin, TX 78712; Howard Hughes Medical Institute, University of California, Berkeley 94720

**Keywords:** Exosome, extracellular vesicle, post-transcriptional modification, RNA-binding protein, RNA-seq, thermostable group II intron reverse transcriptase, Vault RNA, YB-1

## Abstract

RNA is secreted from cells enclosed within extracellular vesicles (EVs). Defining the RNA composition of EVs is challenging due to their co-isolation with contaminants, a lack of knowledge of the mechanisms of RNA sorting into EVs and limitations of conventional RNA-seq methods. Here we present our observations using thermostable group II intron reverse transcriptase sequencing (TGIRT-seq) to characterize the RNA extracted from HEK293T cell EVs isolated by flotation gradient ultracentrifugation and from exosomes containing the tetraspannin CD63 further purified from the gradient fractions by immunoisolation. We found that EV-associated transcripts are dominated by full-length, mature tRNAs and other small non-coding RNAs encapsulated within vesicles. A substantial proportion of the reads mapping to protein-coding genes, long non-coding, and antisense RNAs were due to DNA contamination on the surface of vesicles. Nevertheless, sequences mapping to spliced mRNAs were identified within HEK293T cell EVs and exosomes, among the most abundant being transcripts containing a 5’ terminal oligopyrimidine (5’ TOP) motif. Our results indicate that the RNA-binding protein YBX1, which we showed previously is required for the sorting of selected miRNAs into exosomes, plays a role in the sorting of highly abundant small non-coding RNA species, including tRNAs, Y RNAs, and Vault RNAs. Finally, we obtained evidence for an EV-specific tRNA modification, perhaps indicating a role for post-transcriptional modification in the sorting of some RNA species into EVs. The identification of full-length small non-coding RNAs within EVs suggests a role for EVs in the export and possible intercellular functional transfer of abundant cellular transcripts.

**Statement of Significance:** Cells release vesicles containing selectively packaged cargo, including RNA, into the extracellular environment. Prior studies have identified RNA inside extracellular vesicles (EVs) but, due to limitations of conventional sequencing methods, highly structured and post-transcriptionally modified RNA species were not effectively captured. Using an alternative sequencing approach (TGIRT-seq), we found that EVs contain abundant small non-coding RNA species, including full-length tRNAs and Y RNAs. Using a knockout cell line, we obtained evidence that the RNA-binding protein YBX1 plays a role in sorting small non-coding RNAs into a subpopulation of extracellular vesicles termed exosomes. These experiments expand our understanding of EV-RNA composition and provide insights into how RNA is sorted into EVs for export from the cell.

## Introduction

Metazoan cells grown in culture release extracellular vesicles (EVs) into the surrounding medium and free vesicles can be found in all bodily fluids (1). EVs can be categorized into multiple classes based on their size, shape and presumed membrane origin. Exosomes are defined as ~30-100 nm vesicles that originate from the multivesicular body (MVB) and contain late endosomal markers (2, 3), but there is evidence that biochemically indistinguishable vesicles can bud directly from the plasma membrane (2, 4). Microvesicles or shedding vesicles are generally larger (>200 nm), more variable in shape and density and likely originate from the plasma membrane (1, 3, 5). EVs contain various molecular cargos, including proteins, lipids and RNAs, but how these cargos are sorted into EVs remains unclear.

Since the initial description of RNA in EVs (5-7) and the identification of vesicle-associated circulating transcripts in plasma (8), there has been widespread interest in the possible roles of extracellular RNAs. Seminal EV RNA studies showed the presence of ribonuclease (RNase)-protected mRNA and microRNA (miRNA) species in blood that might be exploited as biomarkers (8) and suggested a role for EVs in the horizontal transfer of genetic information between cells (5-7, 9-11). Several studies implicated miRNAs, in particular, as a mode of intercellular communication based on reporter assays for miRNA function in recipient cells (9-12). Recent studies using sensitive Cre recombinase-based genetic systems in mice have indicated that local and long-range RNA transfer via EVs can occur *in vivo* (13, 14). High-throughput sequencing of extracellular RNA has identified many other EV-associated RNA biotypes, including various small non-coding RNAs (sncRNA, e.g. snoRNA, tRNA, Y RNA, Vault RNA) (15-17), long non-coding RNA (lncRNA) (16, 18) and protein-coding transcripts (16, 19).

The recent application of thermostable group II intron reverse transcriptases (TGIRTs) to sequence low input RNA from human plasma samples revealed that many circulating sncRNA transcripts are full-length and thus apparently resistant to serum ribonucleases (20). In addition to vesicles, extracellular RNA can be found as circulating ribonucleoprotein particles (RNPs), which co-sediment with vesicles during standard EV-isolation protocols culminating in high-speed (>100,000Xg) ultracentifugation (21). The co-isolation of RNPs and multiple EV sub-populations has made it difficult to identify the means by which RNA biotypes are exported and how their transfer between cells might occur.

Understanding the mechanisms by which transcripts are sorted into EVs has proven challenging. Argonaute proteins bind mature miRNAs in the cytoplasm and it has been shown that Argonaute 2 (Ago2)-associated miRNAs can be sorted into exosomes in a process regulated by KRAS signaling (22, 23). We previously used a cell-free reconstitution approach to identify a requirement for the RNA-binding protein (RBP) YBX1 in the sorting of a non-Ago2-bound miRNA (miR-223) into exosomes (24), suggesting an RBP chaperone-mediated sorting pathway. Other RBPs have been shown to sort miRNAs in other cell types. A sumoylated form of hnRNPA2B1 was shown to bind miRNAs containing a tetranucleotide motif (GGAG) that are exported from T-cells (25). SYNCRIP (hnRNPQ) recognizes a GCUG sequence at the 3' end of miRNAs from hepatocytes for sorting into EVs (26). Also in hepatic cells, HuR (ELAVL1) has been shown to control the export of miR-122 during the starvation stress response (27). Despite several examples of RBP-mediated sorting, it is not clear if RBPs play a broad role in defining the RNA composition of EVs or package only a select few transcripts. Nor is it clear to what extent RBPs play an active role in RNA sorting or are simply present in RNPs that are passively captured by membrane invagination processes that generate EVs. Indeed, to date there has been no description of the role that RBPs play in broadly determining the RNA content of EVs.

Here we present our observations from sequencing RNAs extracted from sucrose density gradient-purified EVs and further purified, CD63-immunoisolated exosomes from HEK293T cells using TGIRT-seq. We found that sncRNAs, including tRNAs, dominated EV-RNA composition. TGIRT-seq revealed that most of the tRNAs were full-length, mature transcripts, rather than fragments as previously reported and contain a novel post-transcriptional modification that is not prevalent in cytoplasmic tRNAs. Comparison of the RNA content of exosomes secreted by wild-type and YBX1-null cells indicated that YBX1 plays a role in the export of abundant sncRNA species, suggesting that RNA-binding proteins play a primary role in defining the total RNA composition of exosomes.

## Results

### Most extracellular vesicle-associated transcripts are encapsulated within EVs

Extracellular vesicles encapsulate RNAs rendering them resistant to degradation by exogenous ribonucleases (7-9). However, prior studies have focused primarily on the RNase protection of individual transcripts rather than the global RNA content. To broadly define the RNA composition of EVs, we employed TGIRT-seq in combination with RNase protection assays in the presence or absence of a detergent to distinguish transcripts inside or outside of membrane vesicles. EVs were isolated from HEK293T cells by differential centrifugation followed by flotation gradient ultracentrifugation into a discontinuous sucrose gradient to purify vesicles away from RNPs (Fig. 1*A*). After flotation, EVs were harvested from the 20/40% sucrose interface and either left untreated or treated with protease and/or RNase in the absence or presence of non-ionic detergent (1% Triton X-100; TX-100). Bioanalyzer traces of the RNA after each condition showed a predominant peak of 60-70 nt, which was resistant to RNase under all treatment conditions except in the presence of detergent (Fig. 1*B*). These results corroborate previous studies showing that the predominant RNAs in EVs are ~60-70 nt (9, 16). However, previous RNA-seq experiments failed to identify the high abundance transcripts corresponding to this size distribution, indicating that conventional RNA-seq methodology does not efficiently capture these species.

Since the retroviral reverse transcriptases (RTs) used routinely to generate RNA-seq libraries are unable to include species with significant secondary structure and are impeded by certain post-transcriptional modifications, we utilized a TGIRT enzyme (TGIRT-III (InGex), a derivative of GsI-IIC RT) to generate cellular and EV libraries for Illumina sequencing (20, 28). The resulting TGIRT-seq datasets for the RNase-protection experiments were denoted EV1-6 (Table S1). Scatter plots based on these datasets showed that treatment of the EV fraction with protease and/or RNase in the absence of detergent had little effect on the relative abundance of different transcripts (Fig. 1*C* and Fig. S1). In contrast, upon the addition of detergent, the relative abundance of most transcripts was substantially decreased by RNase treatment (Fig. 1*D*).Pairwise comparisons showed little change under most treatment conditions (Pearson's coefficient (*r*) > 0.92), except when detergent was present (untreated vs. detergent only; *r* = 0.86) and especially when detergent was added together with RNase (*r* = 0.49-0.70) (Fig. *S1*). The decreased correlation between untreated samples and those treated with detergent alone indicated the presence of endogenous RNases in the EV fractions.

**Fig. 1.**
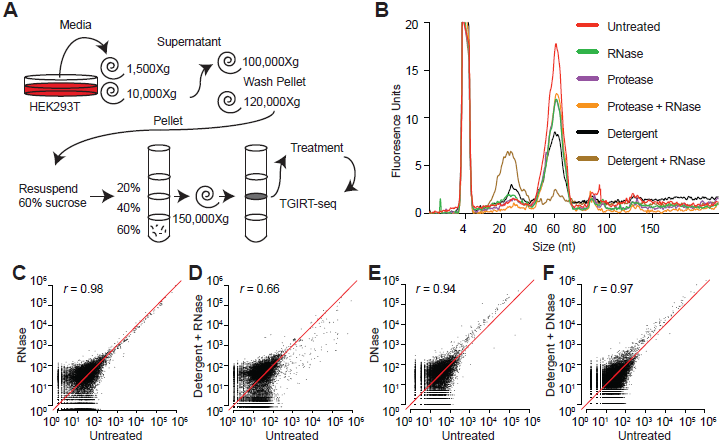
Extracellular vesicle-associated transcripts before and after treatment of EVs with nucleases and protease in the absence or presence of detergent. (*A*) Schematic of the workflow for EV isolation, enzyme and detergent treatment, and TGIRT-seq. (*B*) Bioanalyzer traces (small RNA chip) showing the size distribution of RNAs associated with the EVs before and after the indicated treatments. (*C-F*) Scatter plots comparing DESeq2 normalized read counts between untreated EVs (no detergent and no nuclease) and those treated with RNase or DNase in the absence or presence of detergent. Correlations were computed as Pearson’s correlation coefficients (*r*).

To assess DNA contamination in the EV-RNA isolation, the same treatment conditions used for RNase protection were replicated for DNase protection using a different batch of EVs (datasets EVD1-6; Table S1). Because the TGIRT enzyme can copy DNA fragments efficiently (20, 29), it was possible that some of the sequencing reads were derived from contaminating DNA. As expected, there was no significant effect of detergent or protease treatment on the relative abundance of the large majority of transcripts (r >0.96), and only a minor effect in some datasets when DNase was added in the presence or absence of detergent (r = 0.94-0.97; Fig. 1*E,F*, and r = 0.84-0.97 in the larger set of pairwise comparisons; Fig. S2). Experiments below suggested that the latter reflects cellular DNA that adheres to the surface of the vesicles, even after flotation gradient centrifugation. Together, these results demonstrate that the majority of transcripts recovered from a purification employing a flotation gradient ultracentrifugation step are encapsulated within extracellular vesicles.

### The most abundant transcripts in EVs are full-length tRNAs and other small non-coding RNAs

Analysis of the TGIRT-seq datasets revealed that the most abundant transcripts in the untreated EVs from the RNase-protection experiments were sncRNAs; 62%), followed by ribosomal RNA (27%) and reads mapping to protein-coding genes (6.6%), with the remaining categories accounting for just 3.3% (Fig. 2*A*). Among sncRNAs, by far the most abundant species were tRNAs (50 and 80% of the total and sncRNA reads, respectively) followed by Y RNAs (8.6 and 14% of the total and sncRNA reads, respectively; Fig. 2*A*). Pie charts that were essentially identical to those for the untreated sample were obtained for EVs treated with RNase or with protease and RNase in the absence of detergent, suggesting that there was no substantial RNA or RNP presence on the outside of EVs (Fig. S3). TGIRT-seq revealed only low levels of miRNAs in EVs compared to other transcripts (<0.5% of total reads). Due to secondary structure and post-transcriptional modifications, tRNAs and other sncRNAs may have been under-represented in previous EV small RNA-seq libraries generated by retroviral RTs, thus causing miRNAs to be over-represented, and previous studies often used extraction methods best suited to small RNAs and/or size selection that enriched for miRNAs. Additionally, the version of TGIRT-seq used here relies on a bead-based library clean up that enabled minimal RNA inputs but may have resulted in under-representation of miRNAs, which are close in size to adapter dimers.

**Fig. 2.**
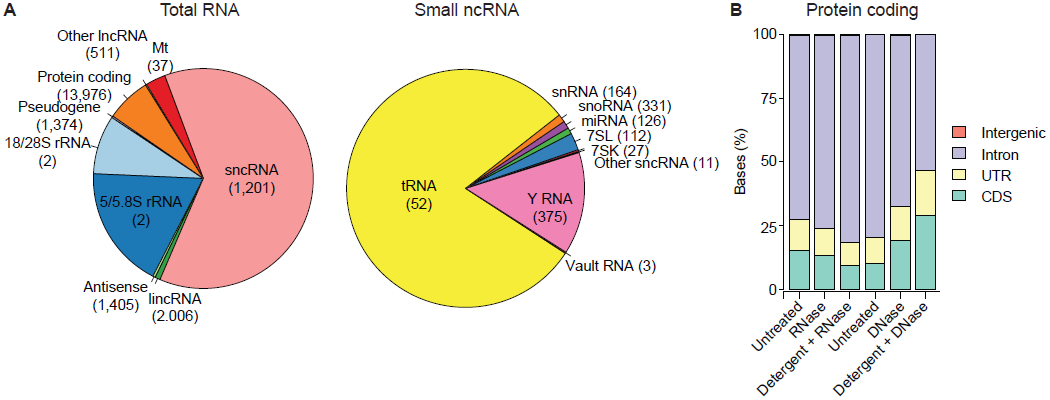
RNA biotypes associated with EVs and the effect of nuclease and detergent treatment on reads mapping to different regions of protein-coding genes. (*A*) Pie charts of percentage of total RNA and small non-coding RNA reads (*left* and *right*, respectively) mapping to the indicated features for untreated EVs. The number of genes represented for each RNA biotype is shown in parenthesis. tRNA gene counts are for tRNA genes grouped by anticodon (N=52, includes three suppressor tRNAs that potentially recognize stop codons), and sncRNA gene counts include pseudogenes. Pie charts of RNA biotypes within the same EVs after RNase or Protease + RNase treatment are shown in Fig. S3. (*C*) Stacked bar graphs of the percentage of protein-coding gene reads mapping to coding sequences (CDS), untranslated regions (UTR), introns, and intergenic regions before and after different treatments. Different batches of EVs were used for the RNase and DNase protection experiments (*left* three bars and *right* three bars, respectively).

Read-span analysis before and after different treatments and Integrative Genomics Viewer (IGV) coverage plots and alignments showed that most of the tRNAs and other sncRNAs present within EVs were full-length, mature transcripts (Fig. 3*A*, Fig. S4, and Fig. S5). In the case of tRNAs, the most abundant RNA species in EVs, read-span analysis showed two prominent peaks, one corresponding to full-length tRNAs and the other to 5’-truncated reads beginning in the D-loop (denoted tRNA*), which results below suggested corresponds to a novel tRNA modification that impedes reverse transcription by GsI-IIC RT. IGV alignments showed that the full-length tRNA reads began precisely at the processed 5’ end of the mature tRNA (or with an extra non-coded G residue in the case of histidyl tRNA), had post-transcriptonal modifications expected for mature tRNAs, and in most cases ended with the post-transcriptionally added 3’ CCA (Fig. S5). Both the full-length tRNAs and tRNA* species were sensitive to RNase in the presence but not absence of detergents and insensitive to DNase, confirming that most tRNAs were enclosed within vesicles (Fig. 3*A*).

**Fig. 3.**
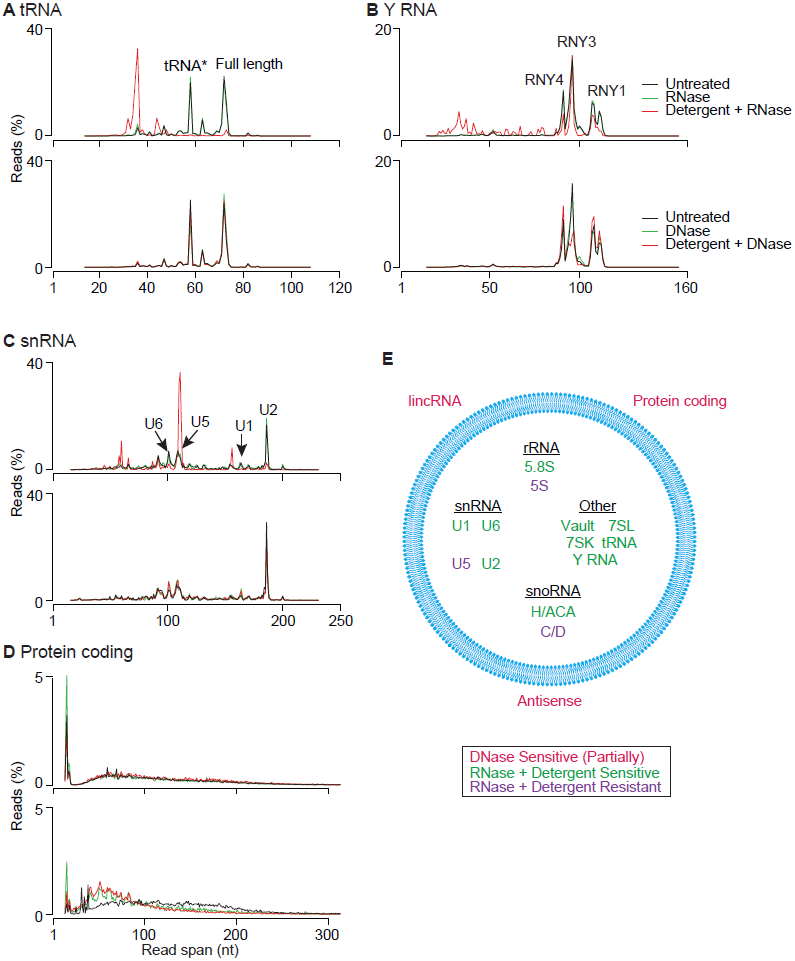
Characterization of EV-RNA biotypes by read span analysis. (*A-D*) Read spans from paired-end sequences of tRNA (A), Y RNA (B), small nuclear RNA (snRNA) (C), and reads mapping to protein-coding genes (D) before and after the indicated treatments (key at *upper right*). The plots show the % of total reads of different read spans for the mapped paired-end reads for each biotype. Peaks for different subspecies of Y RNA and snRNA are indicated on the plots. Results for other RNA biotypes (7SL RNA, snoRNA, 7SK RNA, Vault RNA, 5/5.8S rRNA and 18/28S rRNA) are shown in Fig. S4. (*E*) Summary schematic showing the identity and location of each biotype inside or outside of EVs (blue circle) inferred from nuclease and detergent sensitivity of TGIRT-seq reads.

In addition to full-length tRNAs, read-span analysis and IGV alignments showed the presence of full-length Y RNAs 1, 3, and 4, 7SL and 7SK RNA, both H/ACA- and C/D-box snoRNAs, all species of snRNA, and three species of Vault RNA, whereas only short read spans were found for 18S and 28S rRNA, as expected for Illumina sequencing (Fig. 3, Fig. S4 and Fig. S5). The presence of nuclear snRNAs and 7SK RNA and nucleolar snoRNAs within cytoplasmically derived EVs parallels previous observations for RNAs associated with the cytoplasmic interferon-induced innate immunity protein IFIT5 and host cell RNAs packaged in retroviral virions and may reflect aberrantly localized or non-functional nuclear RNAs that are not associated with their normal partner proteins (30-32). Most species of sncRNAs were efficiently degraded by RNase in the presence of detergent, but several (RNY3, Vault 1-1, C/D-box snoRNAs, 5S rRNA, and U5 snRNA) were resistant to RNase, raising the possibility that they were present in vesicles as RNPs. All of these sncRNA reads were resistant to DNase in the present or absence of detergent as expected (Fig. 3 and Fig. S4).

Our nuclease sensitivity experiments identified reads mapping to protein-coding genes (6.6-10% in the untreated controls for the RNase- and DNase-protection experiments). However, most of these reads mapped to introns rather than coding sequences, and the proportion of reads mapping to introns decreased after DNase treatment in the presence or absence of detergent, suggesting that a significant proportion of these reads come from contaminating DNA (Fig. 2*B*). By read-span analysis, the reads mapping to protein-coding genes appeared as a broad distribution from ~50-200 nt which was degraded to shorter fragments after DNase but not RNase treatment (Fig. 3*D*). Further, this DNase sensitivity was observed irrespective of the presence or absence of detergent in the reaction, indicating DNA contamination on the surface of vesicles. This surface DNA contamination also accounted for a substantial proportion of the reads mapping to lincRNAs, antisense RNAs and other lncRNAs (Fig. S6).

In summary, by combining nuclease treatments with read-span analysis, we found that EVs contained predominantly full-length sncRNAs, particularly tRNAs, and were able to categorize EV-encapsulated RNAs into two distinct classes (Fig 3*E)*: those sensitive to RNase in the presence of detergent (full-length tRNAs, 5.8S rRNA, RNY1 and 4, Vault RNAs 1-2 and 1-3, 7SL and 7SK RNA, H/ACA-box snoRNA, and U1, U2 and U6 snRNAs), and those resistant to RNase in the presence of detergent, consistent with protection by bound proteins in RNPs (RNY3, 5S rRNA, C/D-box snoRNA, U5, snRNA, and Vault RNA1-1). Our experiments also indicated previously unrecognized cellular DNA contamination that adhered to the surface of EVs even after flotation gradient centrifugation and contributed a substantial proportion of the reads mapping to protein-coding genes, lncRNAs and antisense RNAs.

### HEK293T cell exosomes contain spliced mRNAs

Because cells release multiple populations of extracellular vesicles, we focused further analysis on a sub-population of EVs that contain the tetraspanin CD63, likely originating from an endosome-derived MVB, the classical definition of an exosome (24, 33). To isolate CD63^+^ vesicles (exosomes), an additional anti-CD63 antibody affinity purification step was included after the flotation gradient centrifugation (Fig. 4*A*). For the experiments below, we isolated whole cell and CD63^+^ exosomal RNAs from the same cultures and treated the RNAs with DNase prior to TGIRT-seq library preparation. Whole cell RNAs were depleted for rRNA and either fragmented prior to RNA-seq for quantitative comparisons of RNA levels or analyzed directly without fragmentation to preserve intact sncRNAs (34) (datasets WC, WCF, and EX; Table S2).

We confirmed that the RNA content of CD63^+^ exosomes was broadly similar to that of that of the total EV population, with TGIRT-seq showing a similar profile of RNA biotypes, including a high proportion of sncRNAs (73%), with the most abundant being tRNAs (66 and 91% of total RNA and sncRNA, respectively) and Y RNAs (4.0 and 5.5% of total RNA and sncRNA, respectively) and only 1.8% of the reads mapping to protein-coding genes (Fig. 4*B*).

**Fig. 4.**
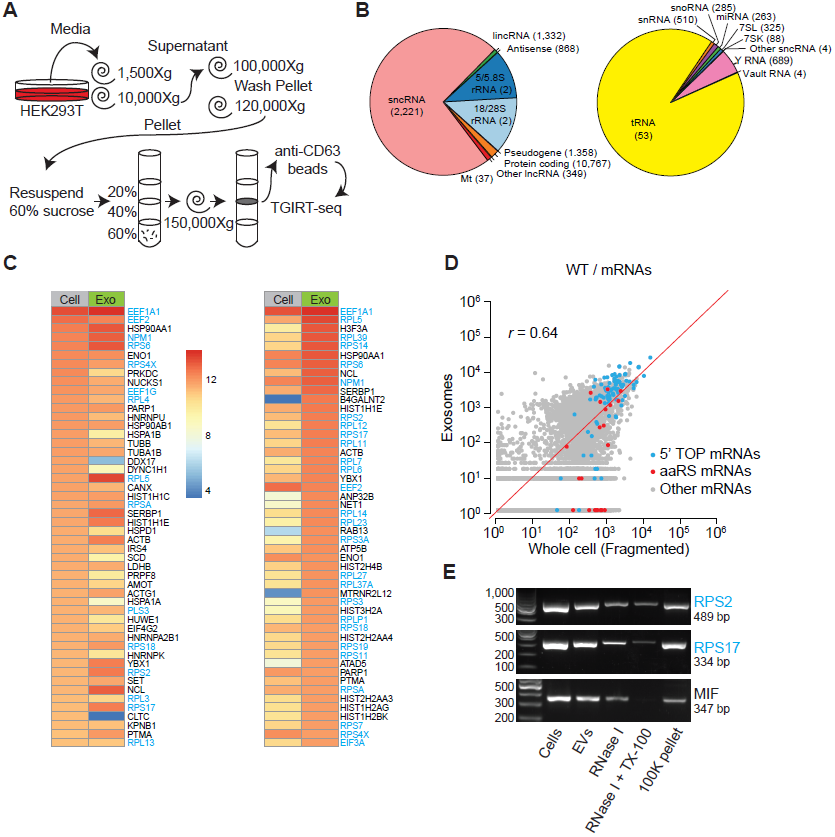
Characterization of RNA biotypes and spliced mRNAs present in CD63^+^ exosomes. (*A*) Exosome (CD63^+^ EV) purification and TGIRT-seq schematic. (*B*) Pie charts showing the percentage of different RNA biotypes in DNase-treated exosomal RNAs from wild-type HEK293T cells. The pie charts show the percentage of total RNA and small non-coding RNA reads (*left* and *right*, respectively) mapping to the indicated features. The number of genes represented for each biotype is shown in parenthesis. tRNA gene counts are for tRNA genes grouped by anticodon (see Fig. 2 legend), and sncRNA gene counts include pseudogenes. (*C, D*) Heat maps (C) and scatter plots (D) showing the relative abundance of different mRNAs in DNase-treated whole cell (fragmented) and exosomal RNAs based on normalized read counts. For this analysis, reads that mapped to protein-coding genes in the initial mapping were re-mapped to the human transcriptome reference sequence (Ensemble GRCh38 Release 76). The heat maps show the 50 most abundant mRNAs by normalized read count in whole cells (left) and exosomes (right), whereas the scatter plots compare all mRNAs. 5' TOP-containing mRNAs, defined as 73 of 80 ribosomal protein mRNAs plus *EEF1A1*, *EEF1B2, EEF1D*, *EEF1G*, *EEF2*, *EIF3A*, *EIF3E*, *EIF3F*, *EIF3H*, *EIF4B*, *HNRNPA1*, *NAP1L1*, *NPM1*, *PABPC1*, *RACK1*, *TPT1*, and *VIM* mRNAs (55), are denoted with blue dots; aminoacyl-tRNA synthetase mRNAs are denoted with red dots. *r* = Pearson’s correlation coefficient. (E) RT-PCR products obtained from RNA isolated from EVs treated with RNase I in the absence or presence of detergent (TX-100). The expected amplicon sizes for *RPS2*, *RPS17* and *MIF* mRNAs are given. The lane at the left shows molecular weight markers.

Our nuclease sensitivity experiments on total EV populations indicated that many of the reads mapping to protein-coding genes reflected DNA contamination on the surface of the vesicles, but left open the possibility that EVs contained smaller amounts of mRNAs or fragments thereof (see above). To address this possibility for CD63^+^ exosomes, we analyzed protein-coding reads that mapped to a human transcriptome reference sequence (Ensembl GRCh38) and compared them to similarly mapped reads for whole cell RNAs from the same cultures (Fig. 4*C*-*E*). In the case of exosomes, the reads mapping to the human transcriptome corresponded to only 0.6% of the total reads, but contained only 0.5-1.2% intron sequences (Fig. S7), indicating that most if not all corresponded to mRNAs or fragments thereof.

Heat maps of the top 50 protein-coding genes showed both similarities and differences between the most abundant mRNA species in cells and exosomes (Fig. 4*C*). Notably, many of the most abundant mRNAs by read count in the whole cell and exosomal RNA preparations encode ribosomal proteins or other proteins involved in translation and belong to a family containing a 5’ terminal oligopyrimidine sequence (5’ TOP), which allows coordinated translational regulation of these transcripts (Fig. 4*C,* gene names highlighted in blue (35-38)).

Scatter plots comparing all putative mRNA species showed some correlation between the relative abundance of different mRNA species in whole cells and exosomes (r = 0.64)(Fig. 4*D*). The majority of the 5’ TOP mRNAs appeared to be over-represented in exosomes, but some were under-represented regardless of whether or not the whole cell RNAs were fragmented prior to sequencing (Fig. 4*D*, Fig. S7B,C). By contrast, reads mapping to mRNAs for aminoacyl-tRNA synthetases (aaRSs), which have previously been reported to be present in exosomes (39), did not appear to be similarly over-represented (Fig. 4*D*, Fig. S7B,C). Since most 5' TOP mRNAs are relatively short (mean transcript length ~ 1 kb), we looked to see if there was a general relationship between transcript length and abundance in exosomes compared to cells. We found that exosomes were enriched in transcripts <1.5 kb, possibly contributing to the enrichment for 5' TOP mRNA sequences (Fig. S8).

To assess whether 5’ TOP mRNA sequences were inside of vesicles, we compared the read coverage across two abundant exosomal mRNAs, *EEF1A* and *RPS2*, after DNase or RNase treatment of EVs in the presence or absence of detergent. We found that reads corresponding to spliced transcripts map across *EEF1A* and *RPS2* exons, unless treated with RNase and detergent, indicating that the mRNAs sequences were enclosed in exosomes (Fig. S9). These results did not distinguish whether these species were full-length mRNA or a series of fragments but minimally demonstrated the presence of multiply spliced mRNA segments inside EVs with an enrichment for 5' TOP-containing transcripts.

To further confirm that spliced mRNAs were present in EVs, we performed reverse transcription PCR using Superscript IV and primers that spanned at least two splice junctions for three putative mRNAs (*RPS2*, *RPL17*, *MIF)*. Spliced mRNAs were detected in exosomes and showed partial sensitivity to RNase alone that was increased in the presence of RNase and detergent (Fig. 4*E*). In summary, we provide evidence for the presence of mRNAs in HEK293T cell exosomes with transcripts containing a 5' TOP sequence motif being particularly abundant.

### YBX1-dependent exosomal RNA transcripts

We previously identified YBX1 as an RNA-binding protein that directs selective miRNA sorting into CD63^+^ exosomes (24). We therefore sought to more comprehensively define the YBX1-dependent exosomal RNA composition by performing TGIRT-seq on DNase-treated RNAs from purified wild-type and YBX1-null cell exosomes (datasets WC, WCF, and WEX for wild type and YC, YCF, and YEX for YBX1-null cells; Table S2). Scatter plots comparing DESeq2-normalized mapped read counts for WT and YBX1-null cellular RNAs did not show dramatic changes in the relative abundance of major exosomal RNA biotypes, except for an increase in moderately expressed tRNAs (Fig. 5*A, left panel*). In contrast, exosomes from YBX1-null cells showed substantially decreased abundance of the large majority of tRNAs, Y RNAs, and Vault RNAs (Fig. 5*A*, *right panel*), whereas other RNA biotypes were not significantly affected (shown by the composite scatter plot in Fig. 5*A*, *right panel* and by separate scatterplots for different RNA biotypes in Fig. S10). Bar graphs of normalized read counts showed that tRNAs were 69% decreased, Y RNAs were 71% decreased, and Vault RNAs were 77% decreased in YBX1-null exosomes (Fig. 5*B*). Conversely, YBX1-null cells showed an increase in the proportion of reads that mapped to tRNAs (2.3-fold), Y RNAs (2.5-fold) and Vault RNAs (1.2-fold) (Fig. 5*B*), suggesting that blocking cellular export via the exosome pathway resulted in accumulation of these transcripts in cells.

**Fig. 5.**
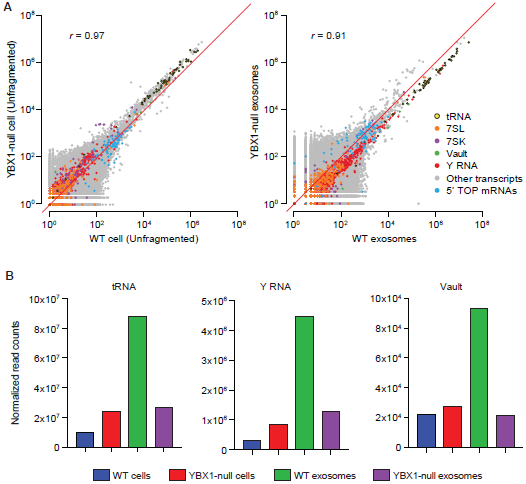
Sequencing of cellular and CD63^+^ exosomal RNA from wild type and YBX1-null cells. (*A*) Scatter plots comparing DESeq2-normalized mapped read counts for the indicated biotypes for DNase-treated WT and YBX1-null cellular RNA (unfragmented; left) and exosomal RNA (right). (*B*) Bar graph showing total normalized read counts mapping to tRNA, Y RNA, and Vault RNA in each library. Normalized read counts represent DESeq2 normalized read counts summed for all transcripts annotated for each indicated biotype in the GENCODE gene set. Bar graphs showing normalized read counts for different subspecies of Y and Vault RNA are shown in Fig. S11.

Further analysis of tRNAs showed that the relative abundance of reads mapping to nearly all tRNA genes were decreased by 50% or more in YBX1-null exosomal RNAs, with most decreased by 70-90% (mean proportion KO/WT = 0.31 (Fig. 6*A*). Individual tRNA isoacceptor levels in cells were increased ~2-fold (mean proportion KO/WT = 2.34) (Fig. 6*A*), further indicating that blocking YBX1-dependent secretion of tRNAs resulted in tRNA accumulation in cells. Thus, the substantial decrease in the proportion of exosomal reads mapping to tRNAs is not confined to a small group of highly expressed tRNA genes. Similarly, different subspecies of Y RNA (RNY1, 3 and 4) and Vault RNA (1-1, 1-2, and 1-3) all showed parallel decreases in YBX1-null exosomes (Fig. S11). Similar decreases in the relative abundance of tRNAs, Y RNAs and Vault RNAs in YBX1-null exosomes were also observed in two earlier versions of this experiment using non-DNase-treated exosomal RNAs. Together, these results suggest a broad role for YBX1 in packaging tRNA as well as some other sncRNA species.

**Fig. 6.**
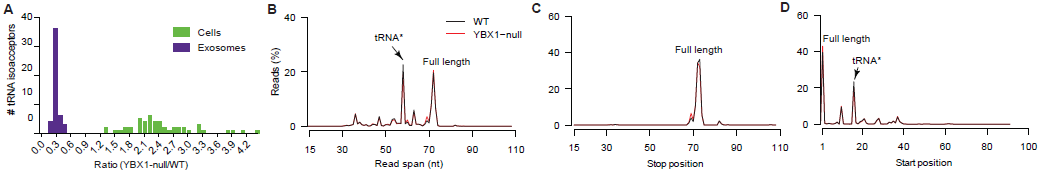
Effect of the YBX1-null mutation on the sorting of different tRNA species into exosomes. (*A*) Histogram showing the number of tRNA species grouped by anticodon (N=49) with the indicated ratio of normalized reads in DNase-treated YBX1-null versus wild-type exosomal RNA preparations. (*B-D*) read span (B), stop site (C) and start site (D) distributions for tRNAs mapped reads as a percentage of total tRNA mapped reads. The peak representing the tRNA* species with a strong stop at position 16 in the D-loop is indicated (B, D).

Read-span analysis showed little difference in the size distribution of any of these sncRNAs between wild-type and YBX1-null exosomes (Fig. S12). In both wild-type and YBX1 exosomes, a high proportion (35%) of tRNA reads spanned the full-length mature tRNA sequence (~75 nt; Fig. 6*B*). Most of the tRNA reads began at the 5’ end of the mature tRNA (position 1), and nearly all tRNA reads terminated at the 3' end of the mature tRNA (positions 72-75) (Fig. 6*C*). The remaining peaks present in Fig. 6*B*, represented 5' truncated tRNA reads, the most prominent being the tRNA* species with a reverse transcription stop at position 16 in the D-loop (Fig. 6*D*).

In contrast to sncRNAs, exosomes from YBX1-null cells did not show a decrease in 5’ TOP mRNAs compared to their levels in cells (Fig. 5*A*, blue dots, compare left and right hand panel). Therefore, YBX1 plays a specific role in defining RNA content of exosomes by influencing the packaging of tRNAs and other sncRNAs.

### An exosome-specific tRNA modification

The results above showed that in addition to full-length tRNAs, EVs and exosomes contained a 15-nt shorter species (tRNA*), which could reflect a D-loop truncation or a post-transcriptional modification that caused a reverse transcription stop. This D-loop stop was not universal for all exosomal tRNAs with ArgCCT, LysCTT, LysTTT, MetCAT, PheGAA, and ValTAC as the primary contributors to this peak (Fig. S13). TGIRT enzymes have been previously shown to read through transcripts with stable secondary structure and post-transcriptional modification (20, 28) and do not stop at the known D-loop modification dihydrouridine, which is prevalent in all tRNAs. Nevertheless, it remained possible that exosomal tRNAs contain a previously unknown modification that results in a more stringent reverse transcription stop.

Since TGIRT enzymes show differential ability to read through modified transcripts, we purified EVs from conditioned medium (according to the protocol in Fig. 1*A*) and split them into two reverse transcription reactions using either GSI-IIC RT (the enzyme used for the previous analysis in this study) or an alternative TGIRT enzyme, TeI4c RT (20, 28) (datasets GC and GEV for GsI-IIC RT and TC and TEV for TeI4c RT; Table S3). The GSI-IIC enzyme gave a read profile with a tRNA* peak in EVs that was greatly diminished in cellular tRNAs from the same culture, demonstrating that the D-loop stop is EV-specific (Fig. 7*A*, left panel). In contrast, TeI4c RT showed only a very small peak at the position corresponding to the tRNA* peak in the same EV RNA preparation and no peak in the same whole cell RNA preparation (Fig. 7*A*, right panel). IGV alignments confirmed that the tRNA* species had a strong reverse transcription stop at the dihydrouridine site for the majority of reads obtained for EV tRNAs with GsI-IIC RT but not with TeI4c RT and was not abundant in reads obtained for cellular tRNAs with both enzymes (shown for LysCTT and ValTAC in Fig 7*B* and Fig. S14, respectively). Because the tRNA* species was highly enriched in EV compared to cellular RNAs, it likely represented an as yet unidentified additional or alternative modification at the dihydrouridine site.

**Fig. 7.**
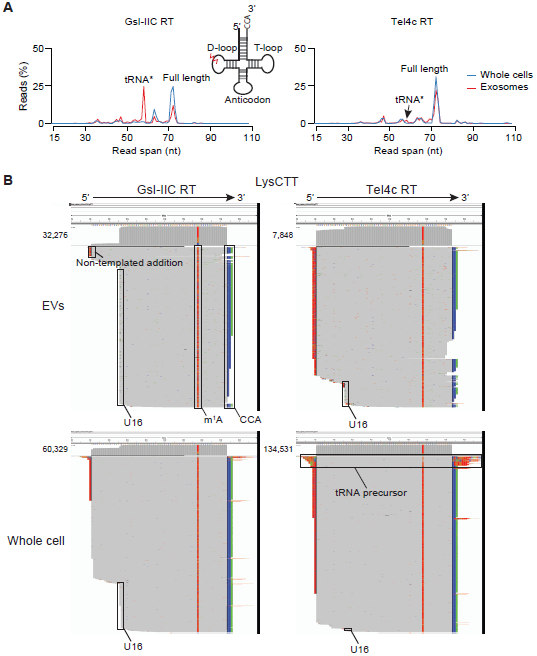
Evidence for an exosome-specific tRNA modification. (*A*) Read span distributions for all reads mapping to tRNAs in TGIRT-seq libraries prepared from unfragmented whole cell RNA and exosomal RNA with either GSI-IIC RT (TGIRT-III; *left*) or TeI4c RT (*right*). The peaks corresponding to full-length mature tRNAs and the tRNA* species with a strong stop at position 16 in the D-loop are indicated. The inset shows a tRNA schematic indicating the approximate position of the putative exosome-specific tRNA modification (jagged red line). (B) Integrative Genomics Viewer (IGV) screen shots showing read coverage across the tRNA coding sequence for a tRNA lysine isoacceptor (CTT) from TGIRT-seq of EV and unfragmented whole cell RNAs prepared with GsI-IIC RT or TeI4c RT. Coverage plots are above read alignments to the tRNA coding sequence with reads sorted by top strand position. The numbers of reads are indicated to the left of the coverage plot and were down sampled to 1,000 reads in IGV for visualization. Nucleotides in reads matching nucleotides in the annotated reference are colored gray. Nucleotides in reads that do not match the annotated reference are color coded by nucleotide (A, green; C, blue; G, brown; and T, red). 1-methyladenine (m1A) and the 3’ CCA post-transcriptional modifications, extra non-templated nucleotides added to the 3’ end of the cDNA by the RT, and the D-loop modification (stop at position U16) are indicated. The reads identified as tRNA precursor have short 5’- and 3’-end extensions that map to the genomic loci encoding this tRNA.

## Discussion

In this report we describe the use of TGIRT-seq, employing a highly processive thermostable group II intron RT, to characterize the RNA content of HEK293T cell EVs and exosomes and to investigate the role of YBX1 in packaging these RNAs. RNase and DNase protection experiments in the presence and absence of detergent show that the most abundant RNA species in both EVs and exosomes are tRNAs and other sncRNA and that a high proportion of these sncRNAs are present as full-length, mature transcripts. tRNAs and most other sncRNA are fully sensitive to RNase in the presence of detergent, but some (5S rRNA, U5 snRNA and some snoRNA, Y RNA and Vault RNA species) are RNase resistant suggesting that they may be protected in RNPs associated with EVs. Results from previous studies from multiple cell types have shown a dominant peak corresponding to full-length tRNA as evaluated by bioanalyzer size distribution (9, 11, 15, 16, 40), but studies which present sequencing results report that the majority of tRNAs are present as fragments (15, 16, 19, 40-42), presumably reflecting the fact that conventional reverse transcriptases arrest at tRNA modification sites. In contrast, a recent study sequencing of cell-free RNA from human plasma by TGIRT-seq revealed the presence of mostly full-length transcripts (20). Our findings using TGIRT-seq clearly demonstrate that EVs contain a high proportion of full-length tRNAs and other sncRNA species (Fig 3 and Fig 7) and indicate that conventional RNA-seq methods are not sufficient to characterize the RNA content of EVs.

The presence of mRNA inside EVs has been controversial in part due to the co-sedimentation of contaminating RNA associated with RNPs present in exosome pellets obtained by high-speed ultracentrifugation or hydrophilic polymer precipitation (*e.g.*, PEG, ExoQuick). A recent report using a multi-stage EV isolation procedure including density gradient flotation identified lincRNA, antisense RNA and protein-coding RNAs as being associated with a particular EV sub-population (40). In contrast, our nuclease sensitivity results indicated that a substantial proportion of the reads mapping to protein coding genes, lincRNAs and antisense RNAs, are due to DNA contamination on the surface of vesicles (Fig. 2 and Fig. 3). These disparate results underscore the importance of both upstream (*i.e.,* EV isolation) and downstream (*i.e*., RNA isolation) purification procedures when characterizing EV composition.

Nevertheless, in addition to sncRNAs, we find smaller amounts of spliced mRNAs (<1 % of total RNA reads) in flotation-gradient purified, DNase-treated RNA from HEK293T cell exosomes (Fig 4). These transcripts are enriched for 5' TOP mRNA sequences (Fig 4*B*, Fig. S7, and Fig. S8). 5' TOP gene products are regulated at the translational level via recruitment to RNA granules in response to starvation (37), perhaps suggesting a link between cellular metabolism and EV-RNA loading. If so, the mRNA content of EVs and exosomes may be cell-type specific and change under different physiological conditions. Our results also confirm a previous report that aaRS mRNA sequences are present in exosomes, of potential interest because of the orthogonal extracellular signaling function of these enzymes (39, 43).

Evidence is accumulating to support a role for RNA-binding proteins in sorting specific transcripts into EVs derived from various cell types. These studies have primarily focused on the sorting of select miRNAs into EVs derived from multiple cells types (see Introduction). To date there has been no attempt to understand how individual RNA-binding proteins define the total RNA composition of EVs beyond specific miRNAs. YBX1 has been shown to bind an array of coding and non-coding cellular transcripts including directly binding tRNA sequences via its cold-shock domain (44-47) and other RNAs non-specifically via its positively charged, intrinsically disordered C-terminal region (48). We find that HEK293T cells devoid of YBX1 show decreased secretion via CD63^+^ exosomes of various abundant sncRNAs, including tRNAs, Y RNAs and Vault RNAs relative to other RNA biotypes (Fig. 5). YBX1 is involved in the formation of stress granules in cells (47, 49) and many other RNA-binding proteins are associated with RNA granules via intrinsically disordered sequences (50). These observations could link RNA granules to the RNA composition of EVs. Translational repression during the generalized stress response may result in an accumulation of free tRNAs that are bound by YBX1 and exported via exosomes.

Finally, using two distinct TGIRT enzymes we find that the major class of truncated tRNA reads (tRNA*) from our sequencing libraries is likely due to a post-transcriptional modification rather than a 5’ tRNA truncation (Fig. 7). The tRNA* species appears highly enriched in EVs, suggesting a possible role of post-transcriptional modification in the sorting of tRNAs into or specialized processing of tRNAs within EVs. The putative modification appears at a nucleotide position normally occupied by dihydrouridine, but the identity of this modification is currently unknown as is whether it occurs before or after uridine reduction in cells or EVs. Recent evidence suggests that Y RNA fragmentation can occur in EVs, indicating that vesicles may enable specialized cell-free biochemical conversions (51). The low yield of RNA in purified exosomes presents a challenge to the identification of the exosome-specific modification. Nonetheless, that TeI4c RT reads through the site of modification without base misincorporation suggests that the modification does not affect Watson-Crick base pairing and therefore may be a bulky group that is added to the ribose moiety or the non-Watson-Crick face of the nucleotide base.

How might the relationship between YBX1, RNA granules and exosome RNA composition contribute to physiology? YBX1 might bind damaged, non-functional, or excess free tRNAs and other sncRNAs that are not associated with their normal protein partners, as an early step that targets these RNAs for sorting into exosomes. This may be particularly relevant under stress conditions where excess transcripts are not beneficial to individual cells. However, purged RNAs including tRNAs or tRNA fragments that are protected inside EVs may be able to travel to nearby actively growing cells and contribute to or regulate recipient cell metabolism. In this way, YBX1 may play a role in resource sharing and/or signaling between cells within tissue microenvironments. Because YBX1 is over-expressed in many malignancies (52), these processes may be particularly important for the survival and growth of cancer cells in the context of the tumor microenvironment. Direct examination and modulation of the relationship between YBX1 localization, EV-RNA content, RNA granules and growth/stress conditions along with the ability to distinguish full-length tRNAs from tRNA fragments using TGIRT enzymes should help to unravel how these relationships influence cellular physiology in health and disease.

## Methods

### Cell lines and growth conditions

Wild-type HEK293T and YBX1-null cells (24) were cultured in Dulbecco's modified Eagle medium with 10% fetal bovine serum (Thermo Fisher Scientific, Waltham, MA). For EV production, cells were seeded at ~10% confluency in 150-mm CellBIND tissue culture dishes (Corning, Corning NY) containing 30 ml of growth medium and grown to 80% confluency (~48 h). Cells grown for EV production were incubated in exosome-free medium produced by ultracentrifugation at 100,000Xg (28,000 RPM) for 18 h using an SW-28 rotor (Beckman Coulter, Brea, CA) in a LE-80 ultracentrifuge (Beckman Coulter).

### Extracellular vesicle and exosome purification

Conditioned medium (210 ml) was harvested from 80% confluent HEK293T cultured cells. All subsequent manipulations were performed at 4 °C. Cells and large debris were removed by centrifugation in a Sorvall R6+ centrifuge (Thermo Fisher Scientific) at 1,500Xg for 20 min followed by 10,000Xg for 30 min in 500-ml vessels using a fixed angle FIBERlite F14-6X500y rotor (Thermo Fisher Scientific). The supernatant fraction was then passed through a 0.22 μM polystyrene vacuum filter (Corning) and centrifuged at ~100,000Xg (26,500 RPM) for 1.5 h using two SW-28 rotors. The pellet material was resuspended by adding 500 μl of phosphate buffered saline, pH 7.4 (PBS) to the pellet of each tube followed by trituration using a large bore pipette over a 30-min period at 4 °C. The resuspended material was washed with ~5 ml of PBS and centrifuged at ~120,000Xg (36,500 RPM) in an SW-55 rotor (Beckman Coulter). Washed pellet material was then resuspended in 200 μl PBS as in the first centrifugation step and 1 ml of 60% sucrose buffer (20 mM Tris-HCl, pH 7.4, 137 mM NaCl) was added and vortexed to mix the sample evenly. The sucrose concentration in the PBS/sucrose mixture was measured by refractometry and, if necessary, additional 60% sucrose buffer was added until the concentration was >48%. Sucrose buffers (40%, 20% and 0% 1 ml each) were sequentially overlaid on the sample and the tubes were centrifuged at ~150,000Xg (38,500 RPM) for 16 h in an SW-55 rotor. The 20/40% interface fraction was collected and either subjected to nuclease protection assays or, for preparation of CD63^+^ exosomes, diluted 1:5 with phosphate buffered saline (pH 7.4) followed by addition of 1 μg of rabbit polyclonal anti-CD63 H-193 (Santa Cruz Biotechnology, Dallas, TX) per liter of original conditioned medium and mixed by rotation for 2 h at 4 °C. Magvigen protein-A/G conjugated magnetic beads (Nvigen, Sunnyvale, CA) were then added to the exosome/antibody mixture and mixed by rotation for 2 h at 4 °C. Beads with bound exosomes were washed three times in 1 ml PBS and RNA was extracted by using a Direct-Zol RNA mini-prep kit (Zymo Research, Irvine, CA).

### Nuclease-protection experiments

Post-flotation EV fractions from the 20/40% sucrose gradient interface (100 μl) were mixed with Triton X-100 (TX-100; 10 μl 10% in 20 mM Tris-HCl, pH 7.4, 137 mM NaCl) or buffer alone. The mixture was then briefly mixed by vortexing and incubated on ice for 30 min. Proteinase K was added to a final concentration of 10 μg/ml to some samples, incubated on ice for 30 min and inactivated by the addition of phenylmethlysufonyl fluoride (PMSF) (5 mM). RNase If (40 U) and 11 μl of New England Biolabs Buffer 3 were then added and incubated at 30° C for 20 min. Alternatively, Turbo DNase (2 U; Thermo Fisher Scientific) and 11 μl of Turbo DNase buffer were added and the solution was incubated at 37 °C for 30 min. Enzymes were inactivated by the addition of 700 μl of Trizol and RNA extraction was performed with the Direct-Zol RNA mini kit.

### RNA-seq library preparation using thermostable group II intron reverse transcriptase (TGIRT-seq)

TGIRT-seq libraries were prepared from 1-3 ng of EV or CD63^+^-exosomal RNA extracted as described above using the TGIRT total RNA-seq method (20, 28, 34). Where indicated, RNAs were treated with DNase I (Zymo Research, manufacturer’s conditions) prior to library construction. In the first step, TGIRT template-switching reverse transcription reactions were performed with an initial template-primer substrate consisting of a 34-nt RNA oligonucleotide (R2 RNA), which contained an Illumina Read 2 primer-binding site and a 3’-blocking group (C3 Spacer, 3SpC3; IDT), annealed to complementary 35-nt DNA primers (R2R DNAs) that have an equimolar mixture of A, C, G, or T single-nucleotide 3’ overhangs. Reactions were performed in 20 μl of reaction medium containing the RNA, 100 nM template-primer substrate, 1 μM GsI-IIC RT (TGIRT-III; InGex, St. Louis MO) or 2 μM TeI4c RT (TeI4c-ΔEn fusion protein RT (20)), and 1 mM dNTPs (an equimolar mix of dATP, dCTP, dGTP, and dTTP) in 450 mM NaCl, 5 mM MgCl2, 20 mM Tris-HCl, pH 7.5, plus 5 mM dithiothreitol (DTT) for GsI-IIC RT and 1 mM DTT for forTeI4c RT. Reactions were assembled by adding all components, except dNTPs, to a sterile PCR tube containing RNAs with the TGIRT enzyme added last. After pre-incubating at room temperature for 30 min, reactions were initiated by adding dNTPs and incubated for 15 min at 60 °C. cDNA synthesis was terminated by adding 5 M NaOH to a final concentration of 0.25 M followed by incubation at 95 °C for 3 min and finally neutralized with 5 M HCl. The resulting cDNAs were purified with a MinElute Reaction Cleanup Kit (QIAGEN) and ligated at their 3’ ends to a 5’-adenlyated/3’-blocked (C3 spacer, 3SpC3; IDT) adapter containing the complement of an Illumina Read 1 primer binding site (R1R) using Thermostable 5’ AppDNA/RNA Ligase (New England Biolabs) according to the manufacturer’s recommendations. The ligated cDNA products were re-purified with a MinElute column and amplified by PCR with Phusion High-Fidelity DNA polymerase (Thermo Fisher Scientific) and 200 nM of Illumina multiplex and 200 nM of barcode primers (a 5’ primer that adds a P5 capture site and a 3’ primer that adds an Illumina sequencing barcode and P7 capture site). PCR was performed with an initial denaturation at 98 °C for 5 sec followed by 12 cycles of 98 °C for 5 sec, 60 °C for 10 sec and 72 °C for 10 sec. The PCR products were purified by using Agencourt AMPure XP beads (Beckman Coulter) to remove primer dimers and sequenced on an Illumina NextSeq 500 instrument to obtain the indicated number of 75-nt paired-end reads or in one experiment on an Illumina HiSeq 2500 to obtained the indicated number of 125-nt paired-end reads (Tables S1 to S3).

The RNA-seq libraries of HEK293 cellular RNAs were constructed similarly. Cellular RNAs (1-1.5 μg) were treated with DNase I (Zymo Research) according to the manufacturer’s protocol, followed by rRNA depletion using a RiboZero™ Gold Kit (Human/Mouse/Rat) (Epicentre). The resulting RNAs (50 ng) were either used directly in the TGIRT template-switching reverse transcription reaction or were fragmented to a size predominantly between 70~100 nt by using an NEBNext® Magnesium Fragmentation Module (New England Biolabs). The fragmented RNAs were then treated with T4 polynucleotide kinase (Epicentre) to remove 3’ phosphates, cleaned up with an RNA Clean & Concentrator™ Kit (Zymo Research) and used for RNA-seq library construction with TGIRT enzymes as described above.

The TGIRT-seq datasets described in this manuscript have been deposited in the National Center for Biotechnology Information Sequence Read Archiv (www.ncbi.nlm.nih.gov/sra eunder accession number SRP108712).

### Bioinformatic analysis

Illumina TruSeq adapters and PCR primer sequences were trimmed from the reads with cutadapt (sequencing quality score cut-off at 20; p-value < 0.01) and reads <15-nt after trimming were discarded. Reads were then mapped with HISAT2 v2.0.2 with default settings to the human genome reference sequence (Ensembl GRCh38 Release 76) combined with additional contigs for 5S and 45S rRNA genes and the *E. coli* genome sequence (Genebank: NC_000913) (denoted Pass 1). The additional contigs for the 5S and 45S rRNA genes included the 2.2-kb 5S rRNA repeats from the 5S rRNA cluster on chromosome 1 (1q42, GeneBank: X12811) and the 43-kb 45S rRNA repeats that contained 5.8S, 18S and 28S rRNAs from clusters on chromosomes 13,14,15,21, and 22 (GeneBank: U13369). Unmapped reads from Pass 1 were re-mapped to Ensembl GRCh38 Release 76 by Bowtie 2 v2.2.6 with local alignment to improve the mapping rate for reads containing post-transcriptionally added 5’ or 3’ nucleotides (*e.g.*, CCA and poly(U)), short untrimmed adapter sequences, or non-templated nucleotides added to the 3’ end of the cDNAs by TGIRT enzymes (denoted Pass 2). The uniquely mapped reads from Passes 1 and 2 were combined by using Samtools. To process multiply mapped reads, we collected up to 10 distinct alignments with the same mapping score and selected the alignment with the shortest distance between the two paired ends (*i.e.*, the shortest read span). In the case of ties between reads mapping to rRNA and non-rRNA sequences, the read was assigned to the rRNA sequence, and in other cases, the read was assigned randomly to one of the tied choices. Uniquely mapped reads and the filtered multiply mapped reads were combined and intersected with gene annotations (Ensembl GRCh38 Release 76) to generate the counts for each individual gene. Coverage of each feature was calculated by Bedtools. To avoid mis-mapping reads with embedded sncRNAs, reads were first intersected with sncRNA annotations and the remaining reads were then intersected with the annotations for protein-coding genes, lincRNAs, antisense, and other lncRNAs. To further improve the mapping rate for tRNAs and rRNAs, we combined reads that were uniquely or multiply mapped to tRNAs or rRNAs in the initial alignments and re-mapped them to tRNA reference sequences (Genomic tRNA Database, and UCSC genome browser website) or rRNA reference sequences (GeneBank: X12811 and U13369) using Bowtie 2 local alignment. Because similar or identical tRNAs with the same anticodon may be multiply mapped to different tRNA loci by Bowtie 2, mapped tRNA reads were combined according to their tRNA anticodon prior to calculating the tRNA distributions. Coverage plots and read alignments were created by using Integrative Genomics Viewer (IGV) (53). Genes with >1,000 mapped reads were down sampled to 1,000 mapped reads in IGV for visualization. For correlation analysis, RNA-seq datasets were normalized for the total number of mapped reads by using DESeq2 (54) and plotted in R. To identify putative mRNAs, we remapped reads that mapped to protein-coding genes in the initial mapping pipeline to the human transcriptome reference (Ensembl GRCh38 releae 76). Reads from different transcript isoforms of the same gene were combined for count normalization in scatter plots. Reads that mapped to putative mRNAs in the human transcriptome reference were retrieved and remapped to the human genome reference using HISAT2 and analyzed by picardtools to calculate the percentage of bases in CDS, UTR, intron, and intergenic regions.

### Reverse transcription PCR

EVs were isolated and left untreated or treated with RNase If and/or detergent as described above. RNA was extracted using the Zymo RNA miniprep kit according to the manufacturer’s instructions. 10 ng of EV or whole cell RNA was reverse transcribed using the SuperScript IV cDNA synthesis kit (Thermo Fisher Scientific) with the supplied oligo(dT) primer. PCR was performed using primers that spanned at least two splice junctions (MIF-F: 5’-TGCCGATGTTCATCGTAAACA, MIF-R: 5’-TTAGGCGAAGGTGGAGTTGT; RPS17-F: 5’-ATAGAAAAGTACTACACGCGC, RPS17-R: 5’-TTAGGCGAAGGTGGAGTTGT; RPS2-F: 5’-TATGCCAGTGCAGAAGCAGA, RPS2-R: 5’-TGGTCAGTGAACTCCTGATA) over 35 cycles (95 °C for 30 sec, 55 °C for 30 sec and 72 °C for 30 sec) with a final extension at 72 °C for 2 min using GoTaq DNA polymerase (Promega). PCR products were separated on a 2.5% agarose gel and visualized using GelRed stain (Biotium Inc.).

## ACKNOWLEDGMENTS

The authors acknowledge the Texas Advanced Computing Center (TACC) at The University of Texas at Austin for providing HPC resources that contributed to the research results reported within this paper. Funding for this work was from the Howard Hughes Medical Institute (R.S.) and National Institutes of Health R01 grant GM37949 (A.M.L.)

## COMPETING INTERESTS

Thermostable group II intron reverse transcriptase (TGIRT) enzymes and methods for their use are the subject of patents and patent applications that have been licensed by the University of Texas and East Tennessee State University to InGex, LLC. A.M.L., some former and present members of the Lambowitz laboratory, and the University of Texas are minority equity holders in InGex, LLC and receive royalty payments from the sale of TGIRT enzymes and kits and from the sublicensing of intellectual property to other companies.

## DATA DEPOSITION

The TGIRT-seq datasets described in this manuscript have been deposited in the National Center for Biotechnology Information Sequence Read Archive (http://www.ncbi.nlm.nih.gov/sra) under accession number SRP108712.

---

**Fig. S1.** RNase protection of EV-associated transcripts. Pairwise scatter plots of DESeq2 normalized read counts and computed Pearson’s correlation coefficients (*upper left* corner) are shown for all RNase-treatment conditions.

**Fig. S2.** DNase protection of EV-associated transcripts. Pairwise scatter plots of DESeq2 normalized read counts and computed Pearson's correlation coefficients (*upper left* corner) are shown for all treatment conditions.

**Fig. S3.** Pie charts of RNAs associated with EVs after treatment with RNase or protease plus RNase in the absence of detergents. (*A,B*) The pie charts show the percentage of total cellular RNA and sncRNA reads (*left* and *right*, respectively) mapping to the indicated features. The number of genes represented for each biotype is shown in parenthesis. tRNA gene counts are for tRNA genes grouped by anticodon (see Fig. 2 legend), and sncRNA gene counts include pseudogenes.

**Fig. S4.** Characterization of additional EV-RNA biotypes by read span analysis. (*A-F*) Read spans from paired-end sequencing for 7SL RNA (A), snoRNA (B), 7SK RNA (C), Vault RNA (D), 5S and 5.8S rRNA (E) and 18 and 28S rRNA before and after the indicated treatment conditions (key at *upper right*). The plots show the % of total reads of different read spans for the mapped paired-end reads for each biotype. Peaks for different subspecies of snoRNA and Vault RNA are indicated on the plots.

**Fig. S5.** Integrative Genomics Viewer (IGV screen shots showing read coverage across the tRNA or sncRNA coding sequence for representative tRNAs and other sncRNAs. The arrow at the top indicates the 5’ to 3’ orientation of the RNA. Coverage plots are above read alignments with reads sorted by top strand position. The numbers of reads are indicated to the left of the coverage plot and were down sampled to 1,000 reads in IGV for visualization. Nucleotides in reads matching nucleotides in the annotated reference are colored gray. Nucleotides in reads that do not match the annotated reference are color coded by nucleotide (A, green; C, blue; G, brown; and T, red) and may correspond to the complementary nucleotide depending on gene orientation. The 1-methyladenine (m1A), m1G, m5C/Cm, non-coded 5’ G residue of HisGTG, and 3’ CCA post-transcriptional tRNA modifications are indicated. The D-loop stop at position U16, a non-coded poly(U) tail at the 3’ end of some Vault 1-1 RNAs, and a previously annotated SNP (dbSNP, NCBI) found in U5 snRNAD-1 are also indicated.

**Fig. S6.** Read-span analysis of (*A*) lincRNAs, (*B*) annotated antisense RNAs and (*C*) other lncRNAs isolated from EVs before and after the indicated treatments (key in panel A). The plots show the % of total reads of different read spans for the mapped paired-end reads for each biotype.

**Fig. S7.** (*A*) Stacked bar graphs of the percentage of reads mapped to the mRNA transcriptome corresponding to coding sequences (CDS), untranslated regions (UTR), introns, and intergenic regions. (*B*) Scatter plots comparing normalized read counts for mRNAs from DNase-treated exosomal RNA and unfragmented whole cell RNAs. (*C*) Scatter plot comparing normalized read counts for mRNAs from DNase-treated unfragmented and fragmented whole cell RNA. 5' TOP-containing transcripts are denoted with blue dots. *r* = Pearson's correlation coefficient.

**Fig. S8.** (*A-D*) Plots showing the normalized read count for mRNAs versus transcript length for exosomal RNA (A), fragmented whole cell RNA (B), unfragmented whole cell RNA (C) and an overlay of whole cell (fragmented) and exosomal RNA (D). (E) Total mRNA mapped reads binned by transcript length for exosome and whole cell RNA. (F) Ratio (exosomes/whole cell RNA) of total mRNA mapped reads for the indicated transcript length bins.

**Fig. S9.** IGV screen shots showing read coverage across the *EEF1A1* and *RPS2* genes in untreated EVs compared to EVs treated with DNase or RNase in the absence or presence of detergent. Coverage plots show the read coverage along the gene body. The number of mapped reads for each condition is indicated to the left of the coverage plot. Read alignments show the mapped reads sorted by genomic coordinates. Nucleotides in reads matching nucleotides in the annotated reference are colored gray. Nucleotides in reads that do not match the annotated reference are color coded by nucleotide (A, green; C, blue; G, brown; and T, red) and correspond to the complementary nucleotide in the mRNA because of the gene orientations in IGV. Splice junctions are depicted in the alignments and thin blue bars between gray reads.

**Fig. S10.** Scatter plots comparing DESeq2-normalized read counts for different RNA biotypes in DNase-treated RNAs isolated from WT and YBX1-null exosomes. sncRNAs include pseudogenes. mRNA counts were generated by collecting reads that mapped to protein-coding genes in the initial mapping and then remapping to the human transcriptome reference sequence (Ensemble GRCh38 Release 76).

**Fig. S11.** (*A, B*) Bar graphs showing normalized read counts mapping to different subspecies of Y RNA (A) and Vault RNA (B) for libraries prepared from DNase-treated WT and YBX1-null whole cell RNA (unfragmented) and exosomal RNA. The bar graphs show DESeq2 normalized read counts summed for all transcripts annotated for each indicated biotype in the GENCODE gene set.

**Fig. S12.** Read-span analysis of sncRNAs in DNase-treated wild-type and YBX-1 null exosomal RNA preparations. (*A-G*) Read spans from paired-end sequencing of tRNA (A), Y RNA (B), 7SL RNA (C), snoRNA (D), snRNA (E), 7SK RNA (F), and Vault RNA (G). The plots show the % of total reads of different read spans for the mapped paired-end reads for each biotype. Peaks for different subspecies of Y RNA, snoRNA, snRNA and Vault RNA are indicated in the plots.

**Fig. S13.** Bar graphs showing the percentage of tRNA* with a stop at position 16 in the D-loop for each tRNA isoacceptor species (N=49) in DNase-treated wild-type and YBX1-null whole cell and exosomal RNAs. The percentages are the proportion of reads for each tRNA species that start between positions 15 and17.

**Fig. S14.** IGV screen shots showing read coverage across the tRNA coding sequence for valine isoacceptor TAC from TGIRT-seq of EV and unfragmented whole cell RNAs prepared with GsI-IIC RT or TeI4c RT. Coverage plots are above read alignments to the tRNA coding sequence with reads sorted by top strand position. The numbers of reads are indicated to the left of the coverage plot and were down sampled to 1,000 reads in IGV for visualization. Nucleotides in reads matching nucleotides in the annotated reference are colored gray. Nucleotides in reads that do not match the annotated reference are color coded by nucleotide (A, green; C, blue; G, brown; and T, red). The 1-methyladenine (m1A) and the 3’ CCA post-transcriptional modifications, extra non-templated nucleotides added to the 3’ end of the cDNA by the RT, and the D-loop truncation (U16) are indicated. Reads identified as tRNA precursor have short 5’- and 3’-end extensions that map to the genomic loci encoding this tRNA.

